# Gold nanocluster mediated delivery of siRNA to intact plant cells for efficient gene knockdown

**DOI:** 10.1101/2021.03.17.435890

**Authors:** Huan Zhang, Yuhong Cao, Dawei Xu, Natalie S. Goh, Gozde S. Demirer, Yuan Chen, Markita P. Landry, Peidong Yang

**Author notes:** Current address: Department of Plant Biology and Genome Center, University of California, Davis, 451 Health Sciences Drive, Davis, CA 95616, USA. H.Z. and Y.C. Contributed equally to this work.

## Abstract

RNA interference (RNAi), which involves the delivery of small interfering RNA molecules (siRNA), has been used to validate target genes in plants, to understand and control cellular metabolic pathways, and as a ‘green’ alternative for crop pest tolerance. Conventional siRNA delivery methods such as viruses and *Agrobacterium*-mediated delivery exhibit limitations in host plant species range and their use can result in uncontrolled DNA integration into the plant host genome. Here, we synthesize polyethyleneimine functionalized gold nanoclusters (PEI-AuNCs) to mediate siRNA delivery into intact plant cells and show these constructs enable efficient gene knockdown. We demonstrate that functionalized AuNCs protect siRNA from RNase degradation and are small enough (~2 nm) to bypass the plant cell wall which exhibits a size exclusion limit of 5-20 nm. These AuNCs in turn enable up to 76.5 ± 5.9% GFP mRNA knockdown efficiency with no cellular toxicity. Our data suggest this simple and biocompatible platform for passive delivery of siRNA into intact plant cells could have broad applications in plant biotechnology.

## Introduction

The emergence of RNA interference (RNAi) technologies have enabled rapid and cost-effective genetic manipulation in plant research at the level of the transcriptome for understanding cellular metabolisms^1^, analyzing gene functions^2^, producing improved crop varieties^3^, and protecting crops against pests^4^. One main way to apply RNAi technologies is to deliver an exogenous small interfering RNA (siRNA) into the cytoplasm, posing a significant challenge in plant-related studies^5^. In comparison with animal cells, mature plant cells have a thick multi-layer cellulose polysaccharide cell wall surrounding the cell membrane, which not only provides structural support and protection to plant cells, but also serves as a barrier that limits biomolecule delivery and has prevented translation of other common abiotic delivery methods (i.e. electroporation, heat shocking) from working in plants^6,7^. Currently, viral vectors are commonly used for delivery of siRNA into intact plants, as they allow for strong siRNA expression without relying on plant transformation^8^. However, this method requires construction of cDNA into the viral vectors, and most viruses are host-specific. Similarly, *Agrobacterium*-mediated transformation, the other common method for RNAi in plants, is also limited to certain plant species^9^, and suffers from the additional potential drawback that random DNA integration can occur during *Agrobacterium* transformation, resulting in endogenous gene disruption and constitutive expression of the transgene.

Nanomaterials and nanostructure-mediated nonviral intracellular delivery have enabled numerous advances in animal research and for diverse biomedical applications^10, 11^. More recently, several studies have also shown that nanomaterials can serve as carriers to deliver DNA plasmid, RNA, or proteins into intact plant cells and protoplasts^7, 12–14^. Mitter *et al.* applied clay nanosheets to deliver pathogen-specific double stranded RNA to intact leaves for crop protection against plant viruses with great success^15^. Other studies have shown that single-walled carbon nanotubes can mediate DNA plasmid delivery and transient protein expression in mature plant leaves, and have also enabled highly efficient siRNA delivery and gene silencing in intact plants^16, 17^. Moreover, DNA nanostructures with certain mechanical properties were able to enter plants for efficacious siRNA delivery in mature plants^18, 19^. Conversely, other nanomaterials that are ubiquitously used for delivery in animal systems such as gold nanoparticles and lipid vesicles, have not yet been reported to enable biomolecule delivery in plants, which could be due in part to plant cell wall excluding entry of abiotic particles above the small plant cell wall size exclusion limit of ~5-20 nm^20^.

Gold nanoclusters (AuNCs) have recently shown numerous successful applications in animal research and biomedical applications. Owning to their ultra-small sizes (~2 nm), high biocompatibility, and strong photoluminescence, AuNCs have shown great potential in various applications, including intracellular delivery^21–23^, imaging and energy transfer^24–26^, and diagnosis in animals^21, 27^. Previously, AuNCs were shown to be small enough to pass the bacterial cell wall and are able to interact with bacteria intracellularly to enable photosynthesis with non-photosynthetic bacteria^24^. Notably, AuNCs synthesis is easier and faster than that of prior siRNA nanocarriers used in plant gene silencing applications. Therefore, herein we assess whether AuNCs can pass the plant cell wall to serve as efficient siRNA delivery vehicles. To this end, we synthesized polyethyleneimine functionalized AuNCs (PEI-AuNCs) with three different PEI polymer lengths to enable siRNA loading onto AuNCs. We further quantified the polynucleotide loading capacity of various PEI-AuNCs, tested the DNA-loaded PEI-AuNC conjugate’s ability to internalize into the cytosol of mature plant leaf cells, and quantified the resulting siRNA-mediated gene silencing efficiencies at the mRNA and protein levels. Our results below suggest AuNCs can serve as biocompatible, easily synthesized, and efficient carriers for siRNA delivery and transient gene silencing applications in plants.

## Results and Discussion

### Synthesis and Characterization of PEI modified gold nanoclusters (PEI-AuNCs)

In this work, we synthesized and validated PEI-AuNCs as an effective siRNA delivery and gene silencing platform in mature plants. To synthesize PEI-AuNCs, siRNA was co-incubated and adsorbed onto PEI-AuNCs through electrostatic interactions then delivered *via* abaxial leaf infiltration to silence green fluorescent protein (GFP) transgene expression in transgenic mGFP5 *Nicotiana benthamiana* (*Nb*) plants (Figure 1a).

**Figure 1.**
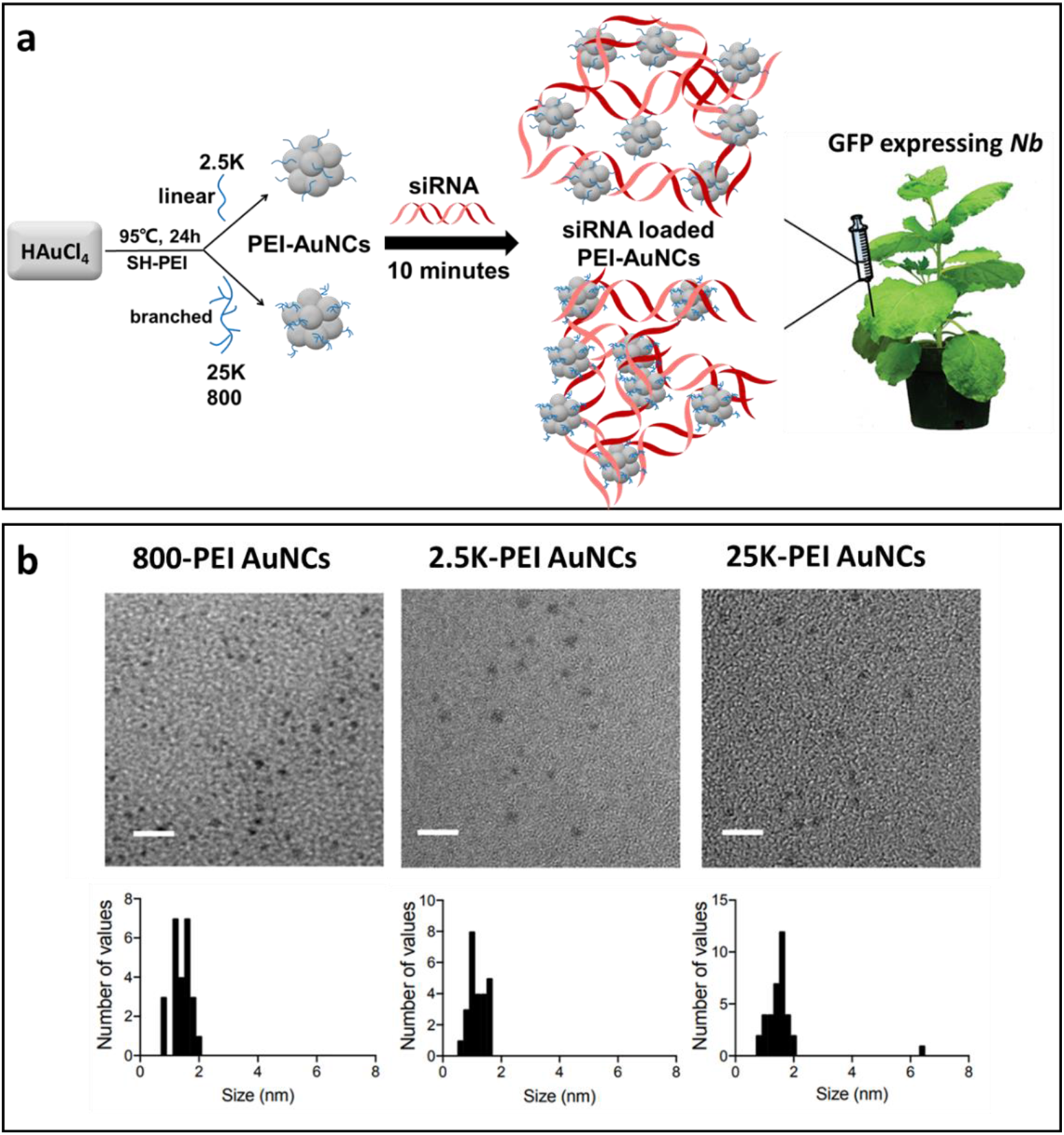
PEI-AuNC synthesis and characterization. **a**, Schematic of PEI-AuNC synthesis (with average PEI molecular weight of 800, 2.5K, and 25K g/mol), followed by siRNA loading through electrostatic adsorption and infiltration-based delivery into mature mGFP5 *Nb* plant leaves for gene silencing. **b**. TEM characterization and size distribution analysis of AuNCs modified by different types of PEI polymers. Scale bar: 10 nm.

We synthesized the PEI-AuNCs with a one-step reduction of gold precursor in the presence of three types of lipoic acid-PEI (~2.5K g/mol linear PEI, ~800 g/mol and ~25K g/mol branched PEI)^28, 29^. Lipoic acid groups are utilized because they can strongly coordinate onto the inorganic surfaces of gold nanocrystals and can be easily combined with tunable length polymers^30^. The 800, 2.5K, and 25K PEI-AuNCs had an average positive Zeta potential of +8.69 mV, +15.97 mV, +20.56 mV, respectively (Table S2), attributed to the PEI ligands that carry amine-derived positive charges. Transmission electron microscopy (TEM) images and corresponding size distribution histograms of the PEI-AuNCs revealed that the 800, 2.5K, and 25K PEI-AuNCs were well dispersed, with well-defined core structures 1-2 nm diameter in size (Figure 1b).

### AuNCs protect loaded siRNA against nuclease degradation

We assessed the siRNA loading ability and gene silencing capacity of the three PEI-AuNCs by incubating different amounts of siRNA with PEI-AuNCs and subsequently assessing the conjugate stability and the siRNA adsorption to the AuNC. The UV-Vis spectra of PEI-AuNCs before and after loading siRNA showed the same characteristic absorption peaks at 522 nm, 371 nm, and 368 nm for 800, 2.5K, and 25K AuNCs respectively, indicating that PEI-AuNCs retain their chemical features and colloidal stability after siRNA loading (Figure 2a). Native PAGE electrophoresis analysis using normalized 120 ng (10 μM, 1 μl) siRNA as a control show that to load 120 ng siRNA, at least 4.8 μg 800 PEI-AuNCs are required, while only 80 ng of 2.5K PEI-AuNCs and 120 ng of 25K-PEI AuNCs are required (Figure 2b, c and d). We attribute the differences in siRNA loading capacity between AuNCs to the inherent differences in size and charge of the constituent PEIs, which may affect the mechanism of interaction between the siRNA cargo and the PEI-AuNC carrier. Based on prior work showing that 100 nM siRNA is optimal for GFP transgene silencing in mGFP5 *Nb*^17^, we thus loaded 100nM siRNA on PEI-AuNCs for our downstream studies. Dynamic light scattering (DLS) measurements revealed that the hydrodynamic size of the PEI-AuNCs increased from 5-7 nm to 20-27 nm after siRNA loading, suggesting adsorption of siRNA to the PEI-AuNCs. Moreover, we observed that siRNA loading on PEI-AuNCs contributes to both increased homogeneity and colloidal stability of the complexes as indicated from the lower polydispersity index (PDI) values and higher zeta potential values following siRNA addition (Table S2).

**Figure 2.**
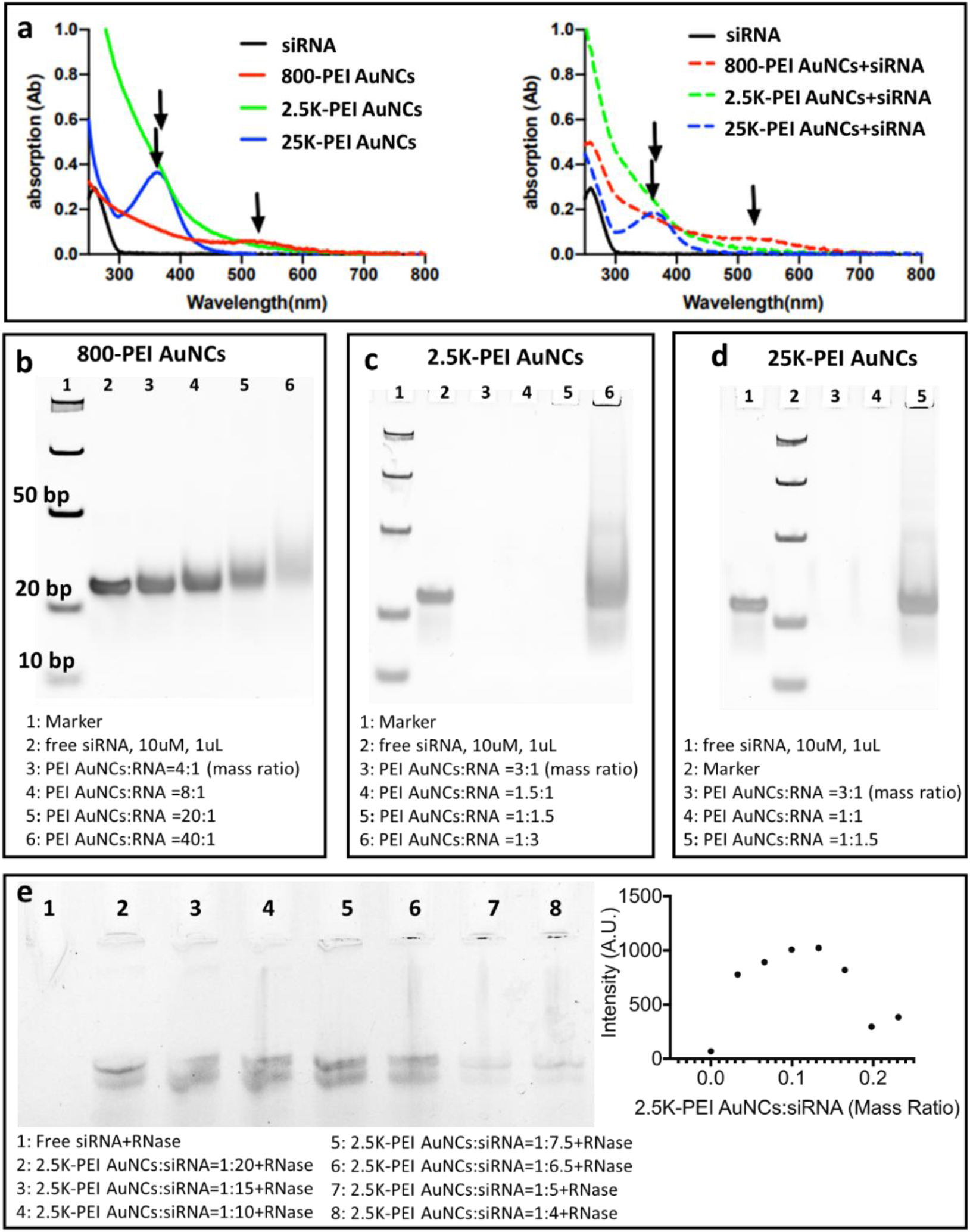
siRNA loading onto AuNCs and protection against nuclease degradation. **a**, Uv-Vis absorbance spectra of PEI-AuNCs before (left) and after (right) siRNA loading (with siRNA only as a control). The corresponding characteristic peaks (black arrows) of different PEI-AuNCs do not shift following siRNA addition, indicating the siRNA-loaded AuNCs remain homogeneous and colloidally stable after siRNA loading. **b-d,** 10% Native PAGE gel to quantify the siRNA loading capacity of the 800-PEI AuNCs (b), 2.5K-PEI AuNCs (c) and 25k-PEI AuNCs (d). **e**, 20% Native PAGE gel showing that siRNA is protected from RNase degradation after being loaded onto 2.5K-PEI AuNCs (the double bands in the gel are caused by bis-intercalating dyes^31^). The right plot is the band intensity analysis of the left gel, where disappearance of the siRNA band represents siRNA degradation.

Since RNA is highly susceptible to degradation, we next investigated the stability of the siRNA once loaded on the PEI-AuNCs. To do so, we incubated either 106 ng free siRNA or siRNA loaded on PEI-AuNCs with RNase H to a final concentration of 10μg/ml at 37°C for 30 min. The reaction products were then run on a 20 % native page gel to quantify the relative amounts of intact versus degraded siRNA. Free siRNA was completely digested as seen from the absence of a siRNA band in lane 1 (Figure 2e). However, when the siRNA was first loaded onto 2.5K-PEI AuNCs, the siRNA was greatly protected as shown by the persistence of an siRNA band across each lane, for a range of siRNA:PEI-AuNC loading ratios (lane 2-6), with a saturating level of PEI-AuNCs observed in lanes 7-8 that retards siRNA migration into the gel. A similar siRNA protection against nuclease degradation is also shown for 800 and 25K PEI-AuNCs (Figure S2).

### Internalization of Cy3 DNA labeled PEI-AuNCs into mature plant cells

We next tested the ability for PEI-modified AuNCs to internalize into the cytosol of mature mGFP *Nb* plant leaf cells. Downstream internalization and silencing experiments were performed with 2.5K-PEI AuNCs since this nanocluster construct was found to have the highest loading capacity (Figure 2a). To do so, we loaded 2.5K-PEI AuNCs with a Cy3-tagged ssDNA oligo. Briefly, 100 ng of Cy3-DNA loaded onto 2.5K linear PEI-AuNCs were introduced into intact mGFP5 *Nb* plant leaves by infiltrating the abaxial surface of the leaf lamina with a needleless syringe (Figure 3a). Following a range of incubation times between 20 minutes and 24 hours post-infiltration, confocal microscopy was performed on the infiltrated leaf tissues. The intrinsic GFP fluorescence of the mGFP5 *Nb* transgenic plant cells provided a fluorescent intracellular marker and thus a metric by which to assess the relative internalization efficiencies of PEI-AuNCs into plant cells. We evaluated the internalization of Cy3 PEI-AuNCs in two ways. First, we changed the mass ratio between PEI-AuNCs and Cy3-DNA from 0 to 100:1 and tested whether the PEI-AuNCs are essential for efficient cellular internalization of DNA. We found that GFP signal highly colocalizes with Cy3 signal when the PEI-AuNCs:DNA ratio is larger than 50:1, suggesting Cy3-DNA requires the PEI-AuNCs carriers for plant cell internalization (Figure 3b). Second, we tested the relative internalization efficiency of Cy3-labeled PEI-AuNCs over a series of incubation time courses from 20 minutes to 24 hours prior to imaging the infiltrated mGFP5 *Nb* leaves. Colocalization of the Cy3 fluorescence (indicating PEI-AuNCs) with the GFP fluorescence (indicating plant cell intracellular space) was calculated to determine the relative extent of PEI-AuNC internalization into the plant cell cytosol. Colocalization analysis suggests that internalization becomes apparent above baseline at 0.5 h with a significantly higher degree of colocalization of 42.3 ± 0.5% than at 20 min (0.33 h) and reaches maximum internalization at 1h post-infiltration with a colocalization fraction of 52.3 ± 2.1% (Figure 3c and d). As the incubation time increases beyond 1h post-infiltration, the colocalization fraction decreases, which might be caused by movement of the PEI-AuNCs out of the cytosol or quenching of the Cy3 fluorescence in the intracellular environment. Representative confocal images in in Figure S1 (all time courses) also support the colocalization trend shown in Figure 3c.

**Figure 3.**
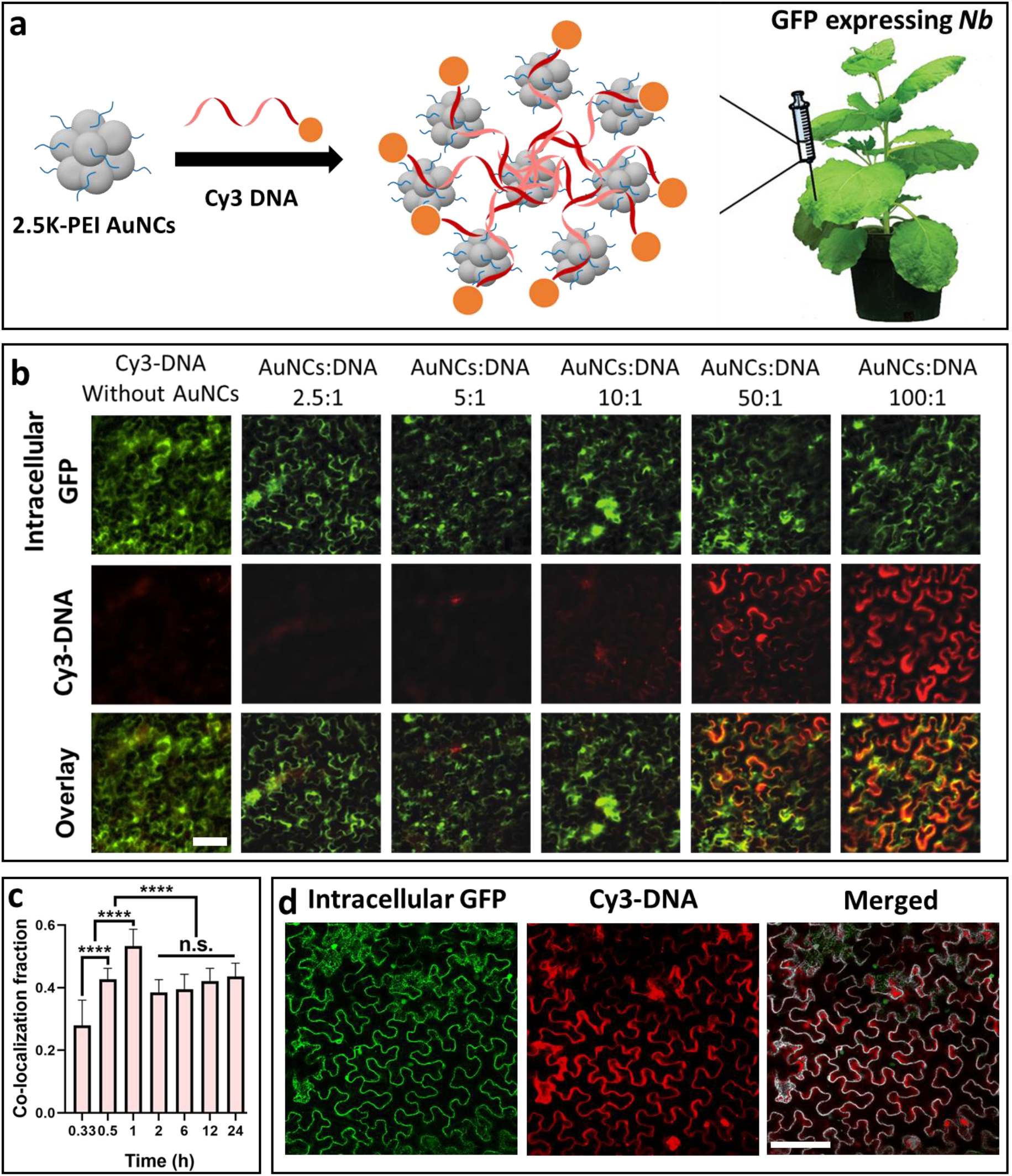
Internalization of Cy3-labeled 2.5K PEI-AuNCs into cells of mGFP5 *Nb* leaves. **a**, Loading of 100 ng of Cy3-labeled single stranded DNA onto 2.5K PEI-AuNCs and infiltration into mGFP5 *Nb* plant leaves. **b**, Confocal images after 1h incubation showing that Cy3-DNA can enter plant cells only if loaded on AuNCs, with most internalization observed at AuNC:DNA mass ratios above 50:1. Scale bar: 100 μm. **c**, Statistical co-localization analysis of Cy3 labelled AuNCs with cytoplasmic GFP at different incubation time points from 20 mins to 24h ****p<0.0001 in one-way ANOVA, n.s.: not significant; Error bars indicate s.e.m. n = 3. **d**, Representative confocal images show internalization of Cy3-DNA loaded AuNCs into mGFP5 plant cells 1-hour post-infiltration. Scale bar: 100 μm.

### PEI-AuNCs mediate siRNA delivery and induce efficient gene silencing in mature plants

Based on the above results suggesting that siRNA can be efficiently loaded onto, protected by, and carried into plant cells by PEI-AuNCs, we tested whether functional siRNA can be delivered by PEI-AuNCs to silence a constitutively expressed GFP transgene in mGFP5 *Nb* plant leaves. We selected a 21-bp siRNA sequence that targets the mGFP5 transgene as a model system to evaluate the ability of PEI-AuNCs to serve as an siRNA delivery tool for transient gene silencing with 800, 2.5K, and 25K PEI polymers. 120 ng (100nM, 100 μl) of this 21-bp siRNA sequence, which inhibits GFP expression in a variety of monocot and dicot plants,^32^ was loaded onto PEI-AuNCs via electrostatic adsorption at a 1:1 mass ratio for 2.5K and 25K PEI-AuNC:siRNA, and a 40:1 mass ratio for 800 PEI-AuNCs for full loading of siRNA on each AuNC. The siRNA-loaded PEI-AuNCs were then infiltrated into the abaxial mGFP5 *Nb* leaf side prior to quantification of GFP transgene silencing 1-day post-infiltration.

To quantify the degree of GFP silencing, we performed quantitative polymerase chain reaction (qPCR) of mRNA transcripts to evaluate siRNA gene knockdown efficiency. mGFP5 *Nb* leaf tissues infiltrated with water, 120 ng free siRNA, 120 ng siRNA loaded onto free PEI polymers, and 120 ng siRNA loaded PEI-AuNCs were quantified with qPCR 1-day post-infiltration. We found that that free siRNA and siRNA mixed with PEI polymers alone do not exhibit any statistically significant decreases in GFP mRNA fold-changes relative to the water-infiltrated leaf control, whereby the siRNA delivered by 25K, 800, and 2.5K PEI-AuNCs show a 63.8 ± 17.8%, 71.2 ± 3.9%, and 76.5 ± 5.9% reduction in GFP mRNA transcript levels, respectively (Figure 4a). We do not observe a statistically significant difference in silencing efficiencies enabled by the different PEI-AuNCs each loaded with 120 ng siRNA, suggesting all three can internalize into plant cells to enable transient suppression of GFP expression. However, we note that the loading efficiency of siRNA onto PEI-AuNCs is highest for 2.5K PEI-AuNCs, and that the 2.5K AuNC clusters therefore represent the best-performing AuNCs for gene silencing on an siRNA mass basis.

**Figure 4.**
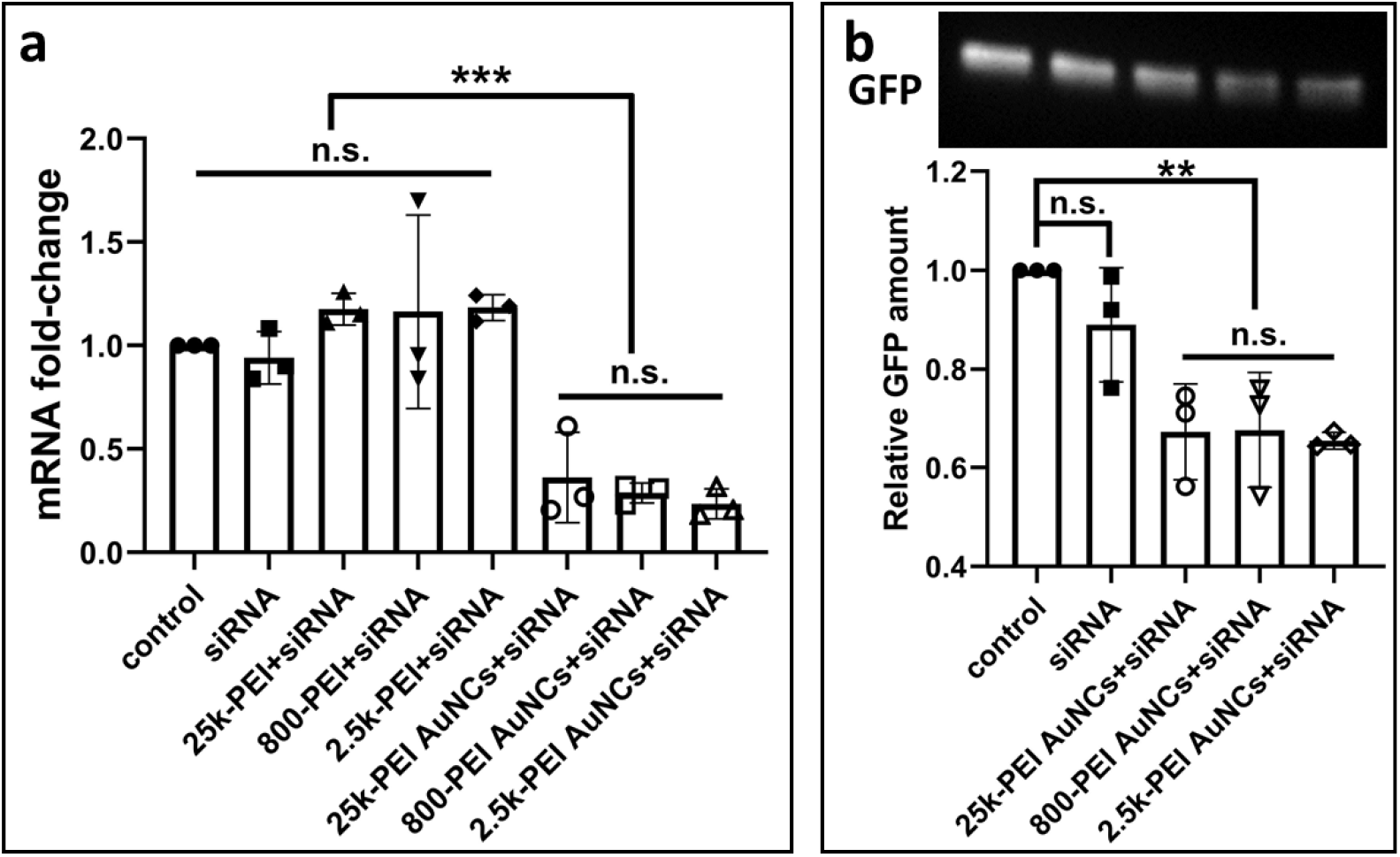
siRNA delivered by 800, 2.5K, and 25K PEI-AuNCs can induce efficacious gene silencing. **a**, qPCR to quantify GFP mRNA-fold changes 1 day post-infiltration with water (control), free siRNA, siRNA mixed with free PEI polymers (800 2.5K and 25K), and siRNA loaded onto PEI-AuNCs. ***P =0.004 in one-way ANOVA. n.s.: not significant; Error bars indicates s.e.m (n = 3). **b**, Representative western blot gel (top image) and statistical analysis of GFP proteins extracted from leaves treated with water (control), free siRNA, or siRNA loaded PEI-AuNCs 3 days post-infiltration. **P = 0.0034, in one-way ANOVA. n.s.: not significant; Error bars indicates s.e.m (n = 3).

To further confirm that PEI-AuNC mediated siRNA intracellular delivery and can silence GFP transgene expression, we quantified GFP levels in leaf tissues 3 days post-infiltration with siRNA-loaded PEI-AuNCs. Treated leaves were excised, proteins were extracted form ~100 mg of treated leaf tissue, and GFP was detected and quantified with western blot analysis. Leaves treated with free siRNA showed the same level of GFP as the water infiltration-treated control, whereas PEI-AuNC treated leaves showed a statistically significant 32-35% reduction in GFP 3-day post infiltration (Figure 4b). Lastly, we tested the biocompatibility of the PEI-AuNCs with qPCR analysis of respiratory burst oxidase homolog B (*NbrbohB*), a ubiquitous plant biotic and abiotic stress gene across many plant species.^33^ Upregulation of *NbrbohB* can be suggestive of stress to plant tissues, and was thus quantified to assess the toxicity of our PEI-AuNCs. As detailed in figure S3, qPCR analysis of *NbrbohB* show that the both the siRNA treated and PEI-AuNCs-treated leaves do not up-regulate *NbrbohB* compared to water-treated control leaves, which suggests that PEI-AuNCs are a biocompatible carrier for siRNA delivery to plants. In sum, PEI-AuNCs yield significantly reduced expression levels of both GFP and GFP mRNA following infiltration into mGFP5 *Nb* leaves and showed unchanged levels of *NbrbohB* stress gene expression. Taken together, our data suggest that PEI-AuNCs can serve as an efficacious platform for siRNA delivery and transient gene silencing in mature plants.

## Conclusion

Gold nanoclusters (AuNCs) can be efficiently, quickly, and reproducibly synthesized to exhibit small size (~2nm) and are amenable to facile surface chemical modification. For these reasons, AuNCs have shown numerous biomedical applications in bioimaging^25, 26, 34^, biomolecule delivery^22, 23, 27^, and bio-sensing^21, 35, 36^. However, to-date, the utility and biocompatibility of AuNCs for use in plants had not been explored. In particular, in contrast to most biological systems, plants exhibit a lower ~20 nm size exclusion limit for biomolecule delivery due to the presence of the plant cell wall. Therefore, AuNCs are promising nanomaterials and potentially suitable for various applications in plant systems due to their inherent ultrasmall sizes and straightforward syntheses. Here, we explored the feasibility of using AuNCs as a carrier core, followed by PEI ligand modification, as an siRNA delivery platform. We demonstrate that siRNA loading capacity varies greatly depending on the nature of the PEI ligand, with 2.5K linear PEI ligands enabling loading of 1.5 μg siRNA per 1 μg AuNCs, relative to the lowest-performing 800 PEI ligands that can only load 25 ng siRNA per 1 μg AuNCs. We further show that PEI-AuNCs can be used as a carrier for efficient transgene gene silencing in mature *Nb* plants, induced by the delivery of siRNA. Characterization of the PEI-AuNCs within the plant leaf cells indicated that compared with other nanomaterials (such as single walled carbon nanotubes, DNA nanostructures), PEI-AuNCs internalize into plant cells relatively quickly (0.5-1h), possibly due to their smaller size. Our data further suggests that the AuNC positive surface charge endowed by PEI ligands enables electrostatic adsorption of polynucleotides and enables siRNA uptake into plant cells. Lastly, we note that the polynucleotide loading mechanism on AuNCs is based on electrostatic adsorption and therefore agnostic to polynucleotide type. Therefore, PEI-AuNCs represent a promising delivery platform for a myriad of plant biotechnology applications, potentially including AuNC-mediated delivery of other DNA or RNA cargoes, e.g. single-guide RNAs and/or mRNAs, or for bioimaging and biosensing applications. In sum, our above results suggest that AuNCs are a promising and biocompatible delivery platform for siRNA mediated gene silencing in mature plants and could provide a broader strategy for diverse plant biotechnology applications.

## Materials and Methods

### Chemicals and materials

Lipoic acid, N, N’-Dicyclohexylcarbodiimide (DCC), Dichloromethane (DCM), Polyethyleneimine (PEI) and Tetrachloroauric (III) acid (HAuCl_4_.3H_2_O) were purchased from Sigma-Aldrich. Slide-A-Lyzer dialysis flasks were purchased from ThermoFisher Scientific. DNAs and RNAs were purchased from Integrated DNA Technologies. RNase was purchased from New England BioLabs.

### Preparation of PEI-lipoic acid

PEI-lipoic acid conjugates were synthesized by coupling lipoic acid to 800, 2.5k and 25k PEI through carbodiimide chemistry as previously reported^37^. In brief, PEI and DCC were first dissolved in ice-cold DCM under nitrogen. After 5 h activation, the reaction mixture was filtered. Then the lipoic acid solution was added dropwise to the filtrate. The mixture was stirred overnight under nitrogen atmosphere at room temperature. The product was purified through ice-cold diethyl ether precipitation and dried in vacuo before use.

### Preparation of PEI functionalized gold nanocluster (PEI-AuNCs)

A solution of 800, 2.5k, and 25k PEI-lipoic acid ligands were prepared in a clear glass vial. To synthesis the PEI-AuNCs, 30 μL of HAuCl_4_·3H2O stock solution (100 mM) was added along with DI water to bring the final volume of the reaction to 6 mL with a molar ratio of ligand to gold of 6:1. Then, 30 μL of 2 M aqueous NaOH solution was added to make the reaction mixture basic. The mixture was heat at 95 °C and left stirring for 24h. An aqueous solution of PEI-AuNCs with yellow color was formed. The PEI-AuNC solution was further purified by dialysis. The purified PEI-AuNCs solution was stored at 4 °C.

### AuNC Characterization

The size distribution, polydispersity index, and surface charge of the prepared nanomaterials were determined using Zetasizer (NanoZS, Malvern). For morphology characterization, the samples were examined by transmission electron microscope (TEM, FEI TECNAI 12 and FEI F20 UT TECNAI). A NanoDrop Microvolume UV-Vis Spectrophotometer (ThermoFisher Scientific) was used to characterize the UV-Vis spectrum of the 800, 2.5k and 25k PEI-AuNCs and siRNA loaded PEI-AuNCs.

### siRNA loading capacity measurements and protection gel assay

Duplex siRNA is obtained by mixing two fully complementary RNA oligonucleotides (sense and antisense RNA strands in Table S1) in PBS buffer and heating to 95°C for 5 minutes, followed by cooling to room temperature (20° C) within 30 minutes. To determine the loading capacity of PEI-AuNCs, pre-hybridized siRNA duplex was incubated with PEI-AuNCs at various mass ratios (4:1, 8:1, 20:1 and 40:1 for 800-PEI AuNCs; 3:1, 1.5:1, 1:1.5 and 1:3 for 2.5K-PEI AuNCs; 3:1, 1:1 and 1:1.5 for 25K-PEI AuNCs) in DI water for 10 minutes. The prepared samples were characterized by 10% native page gel electrophoresis and the band intensity was analyzed and normalized to the free siRNA band (120 ug) to determine how much siRNA was loaded onto the various AuNCs. For the siRNA protection assay, 106 ug of free siRNA or siRNA loaded onto PEI-AuNCs (at various mass ratios of 1:5, 1:15 and 1:30 (PEI-AuNCs:siRNA)) were incubated with RNase H (10μg/ml, New England BioLabs, M0297L) at 37°C for 30 minutes, then the prepared samples were characterized by20% native page gel electrophoresis. The band intensity represents intact RNA, which was analyzed with Image J software.

### RNase Digestion

RNA of 106 ug with different mass ratio of PEI-AuNCs was mixed with 10 mU RNase and incubated at 37° C in a reaction volume of 20 uL for 30 min. After incubation, the mixture was directly load to 20% native PAGE gel for electrophoresis and analysis.

### Plant growth and Maintenance

Transgenic mGFP5 *Nicotiana benthamiana* (*Nb*) seeds (obtained from the Staskawicz Lab, UC Berkeley) were germinated and grown in SunGro Sunshine LC1 Grower soil mixture within a growth chamber (740 FHLED, HiPoint, Taiwan). The plants were grown in 4-inch pots under LED light and a 14/10 light/dark photoperiod at 23°C and 60% humidity. Plants were allowed to mature to 3-4 weeks of age within the chamber before experimental use.

### Leaf Infiltration

A tiny puncture hole was introduced into the *Nb* plant leaf with a pipette tip (10 μl) on the leaf abaxial surface prior to infiltration of leaves. Infiltrations were performed by gently pushing ~100 uL of fluid into the leaf tissue with a 1 mL capacity needle-less syringe.

### Internalization of Cy3 labeled PEI-AuNCs into mGFP5 *Nb* plant leaf cells quantified through colocalization analysis

100 ng of Cy3 modified ssDNA was loaded onto 0.4 ug 2.5K-PEI AuNCs and infiltrated into mGFP5 *Nb* leaves with a final concentration of 200nM Cy3 DNA. The plants were kept in the plant growth chamber for the specified incubation times before preparing infiltrated leaves for confocal imaging. A small leaf section at the infiltration site was cut and mounted between a glass slide and #1 thickness cover slip. Water was added to keep the leaf sections hydrated during imaging. A Zeiss LSM 710 confocal microscope was used to image the plant tissue with 488 nm and 543 nm laser excitation for GFP (collecting window: 490-520nm) and Cy3 (collecting window: 530-600nm) signal collection, respectively. Images were obtained at 20x magnification. The same imaging parameters and quantification analyses were applied to samples imaged at different time points.

### Quantitative PCR (qPCR) experiments for gene silencing

Water (control), free siRNA, siRNA loaded PEI-AuNCs (with total volume of 100 μl and a final concentration of 100 nM siRNA) were infiltrated into plant leaves and left on the benchtop at room temperature (20°C) for 1 day prior to extraction of total RNA of the treated leaves with an RNeasy plant mini kit (QIAGEN). Two-step qPCR was performed to quantify GFP gene silencing using the total RNA extracted from leaves: an iScript cDNA synthesis kit (Bio-Rad) was used to reverse transcribe total RNA into cDNA, and PowerUp SYBR green master mix (Applied Biosystems) was used for qPCR. The target gene in our qPCR was mGFP5 (GFP transgene inserted into *Nb*) as the gene target for RNAi-based silencing, and respiratory burst oxidase homolog B (NbrbohB) was the target gene for toxicity assessments, with EF1 (elongation factor 1) as the invariant housekeeping (reference) gene standard^38^. Primers (see detailed sequences in Table S1) for these genes were ordered from IDT and used without further purification. An annealing temperature of 60°C was used for qPCR, which was run for 40 cycles. qPCR data was analyzed by the ddCt method^39^ to obtain the normalized GFP gene expression-fold change with respect to the EF1 housekeeping gene and control. For each sample, qPCR was performed as 3 technical replicates (3 reactions from the same isolated RNA batch), and the entire experiment consisted of 3 independent infiltrations and RNA extractions from different plants (3 biological replicates).

### Quantitative Western blot experiments and data analysis

Infiltrated plant leaves were harvested 3 days post-infiltration and ground in liquid nitrogen to obtain dry frozen powders. The frozen powders were then transferred to a tube with pre-cooled lysis buffer containing 10 mM Tris/HCl (pH 7.5), 150 mM NaCl, 1 mM EDTA, 0.1% NP-40, 5% glycerol, and 1% cocktail. After lysing on ice for 1h, the tube was centrifuged at 10,000 rpm for 20 minutes and the supernatant containing whole proteins was collected into a new tube. After quantification of the total extracted proteins by a Pierce 660 nm Protein Assay (Thermofisher, 22660), 0.5 μg of normalized total proteins from each sample were analyzed by 12% SDS–PAGE and blotted to a PVDF membrane. The membrane was then blocked for 1 hour using 7.5% BSA in TBST (PBS containing 0.1% Tween 20) buffer and rinsed 3 times in PBST buffer, followed by overnight incubation at 4°C with the primary GFP antibody as required (1:2000 dilution, Abcam, ab290). After extensive washing, the corresponding protein bands were probed with a goat anti-rabbit horseradish peroxidase-conjugated antibody (1:5000 dilution, Abcam, ab205718) for 30 min. After 3 washes with TBST, the membrane was then developed by incubation with chemiluminescence (Amersham ECL prime kit) and imaged with a ChemiDoc™ XRS+ System (BIORAD). The intensity of GFP bands were quantified with Image J software. To correct for variability in protein expression across different plants and leaves, the GFP extracted from each leaf sample was normalized by the total protein recovered from that leaf tissue.

## Acknowledgements

We acknowledge support from a Burroughs Wellcome Fund Career Award at the Scientific Interface (CASI), a Stanley Fahn PDF Junior Faculty Grant under award no. PF-JFA-1760, a Beckman Foundation Young Investigator Award, a USDA AFRI award, a grant from the Gordon and Betty Moore Foundation, a USDA NIFA award, a USDA-BBT EAGER award, support from the Chan-Zuckerberg foundation, and an FFAR New Innovator Award (to M.P.L). Work at the Molecular Foundry was supported by the Office of Science, Office of Basic Energy Sciences, of the U.S. Department of Energy under Contract No. DE-AC02-05CH11231. Authors acknowledge support of the BASF-CARA program. We acknowledge support from Keck Foundation (Grant 89208), and H.Z. acknowledges the support of the start-up fund from Jinan University (88016105). N.S.G is supported by a Foundation for Food and Agriculture Research Fellowship. G.S.D. is supported by a Schlumberger Foundation Faculty for the Future Fellowship. The authors also acknowledge support from UC Berkeley Molecular Imaging Center (supported by the Gordon and Betty Moore Foundation), the QB3 Shared Stem Cell Facility, and the Innovative Genomics Institute (IGI).

## Author Contributions

Yu.C., H.Z., M.P.L. and P.Y. conceived the idea and designed the study. H.Z. and Yu.C. performed the majority of the experiments, data analysis, and wrote the manuscript. D.X. assisted in gold nanocluster synthesis. Yuan.C. helped with plant seeding and maintenance. G.S.D. and N.S.G helped analyze the results. All authors edited the manuscript and approved the final version.

