## Supplemental Information for "Gold nanocluster mediated delivery of siRNA to intact plant cells for efficient gene knockdown"

**Supplementary Table S1.** Sequences of oligonucleotides used in this study.

| Name | Sequence (5’-3’) |
|---|---|
| **Cy3-DNA** | CTAAGGATGCGTGTA-Cy3 |
| **sense RNA** | rGrGrUrGrArUrGrCrArArCrArUrArCrGrGrArATT |
| **antisense RNA** | rUrUrCrCrGrUrArUrGrUrUrGrCrArUrCrArCrC |
| **mGFP forward** | AGTGGAGAGGGTGAAGGTGATG |
| **mGFP reverse** | GCATTGAACACCATAAGAGAAAGTAGTG |
| **EF1 forward** | GCATTGAACACCATAAGAGAAAGTAGTG |
| **EF1 reverse** | ACGCTTGAGATCCTTAACCGCAACATTCTT |
| **NbrbohB forward** | TTTCTCTGAGGTTTGCCAGCCACCACCTAA |
| **NbrbohB reverse** | GCCTTCATGTTGTTGACAATGTCTTTAACA |

**Supplementary Table S2.** Dynamic light scattering (DLS) size and zeta potential measurements of PEI-AuNCs and siRNA loaded PEI-AuNCs.

| Name | Average size (nm) | PDI | Zeta potential (mV) |
|---|---|---|---|
| 800-PEI AuNCs | 6.46 | 0.598 | +8.69 |
| 2.5K-PEI AuNCs | 5.11 | 0.542 | +21.56 |
| 25K-BPEI AuNCs | 6.53 | 0.562 | +15.97 |
| 800-PEI AuNCs+siRNA | 19.58 | 0.353 | +29.50 |
| 2.5K-PEI AuNCs+siRNA | 27.69 | 0.46 | +36.67 |
| 25K-PEI AuNCs+siRNA | 24.42 | 0.382 | +35.7 |

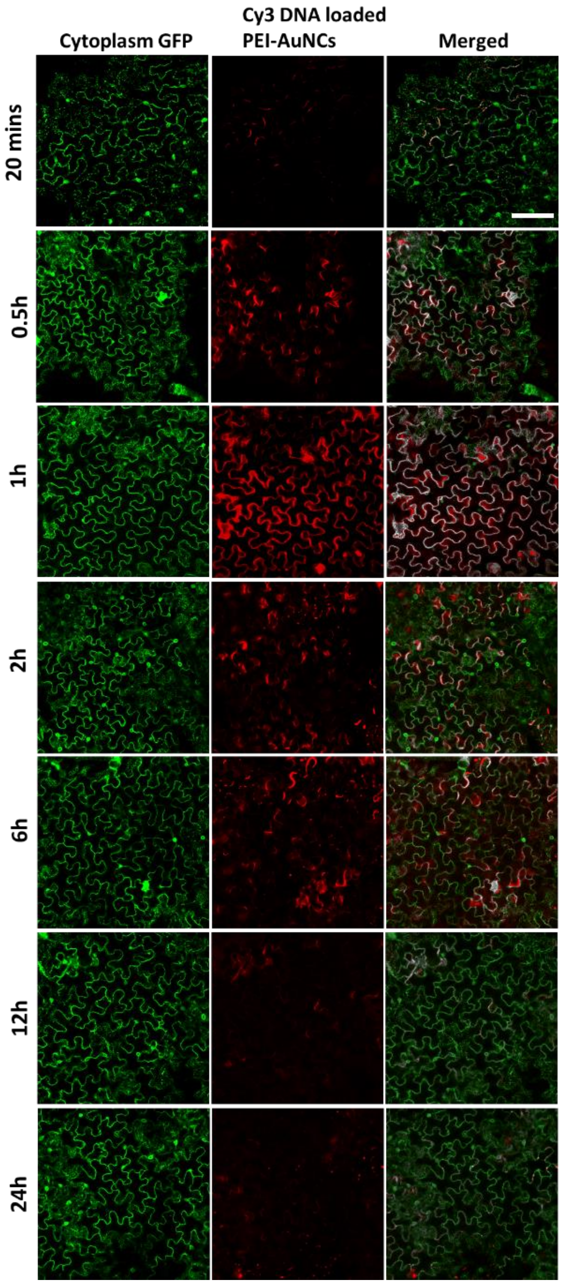


**Figure S1.** Representative confocal images of Cy3 DNA loaded 2.5K-PEI AuNC infiltrated *Nb* leaves at different incubation time points (from 20 mins to 24 h).

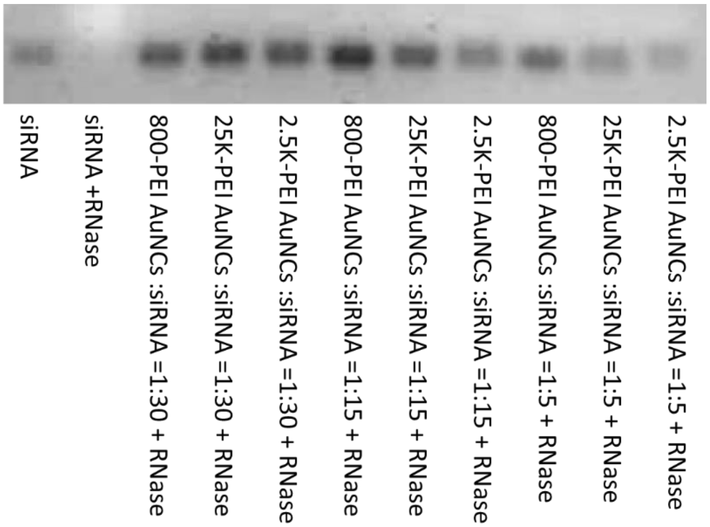


**Figure S2.** 4% Agarose gel indicating protection of siRNA from RNase degradation after loading onto PEI-AuNCs.

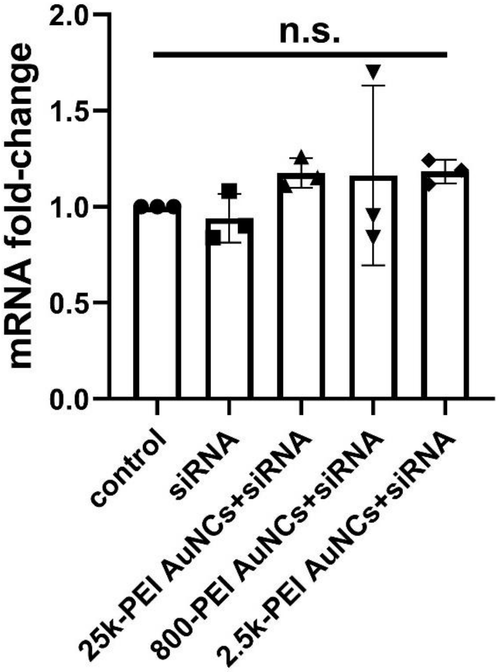


**Figure S3.** qPCR analysis of *NbrbohB* mRNA fold-change1-day post infiltration of samples into *Nb* leaf tissues. n.s.: not significant; Error bars indicates s.e.m (n = 3).
